# MrgprA3-expressing pruriceptors drive pruritogen-induced alloknesis through mechanosensitive Piezo2 channel

**DOI:** 10.1101/2022.06.22.497257

**Authors:** Ping Lu, Yonghui Zhao, Zili Xie, Xinzhong Dong, Gregory F. Wu, Brian S. Kim, Jing Feng, Hongzhen Hu

**Author notes:** These authors contributed equally (Ping Lu, Yonghui Zhao).

## Abstract

Although touch and itch are coded by distinct neuronal populations, light touch also provokes itch in the presence of exogenous pruritogens, resulting in a phenomenon called alloknesis. However, the cellular and molecular mechanisms underlying the initiation of pruritogen-induced mechanical itch sensitization are poorly understood. Here we show that intradermal injections of histamine or chloroquine (CQ) provoke alloknesis through activation of TRPV1- and MrgprA3- expressing prurioceptors, and functional ablation of these neurons reverses pruritogen-induced alloknesis. Moreover, genetic ablation of mechanosensitive Piezo2 channel function from MrgprA3-expressing prurioceptors also dampens pruritogen-induced alloknesis. Mechanistically, histamine and CQ sensitize Piezo2 channel function through activation of the PLC-PKCδ signaling. Collectively, our data uncovered a TRPV1^+^/MrgprA3^+^ prurioceptor-Piezo2 signaling axis in the initiation of pruritogen-induced mechanical itch sensitization in the skin.

## INTRODUCTION

Alloknesis (or mechanical itch sensitization) is defined as an abnormal sensory state where innocuous mechanical stimuli (such as stimuli from clothes) evoke nocuous itch sensation in the settings of pruritogen stimulation and chronic itch. Although the phenomenon is well known clinically, mouse model of pruritogen-induced mechanical itch was not established until the year of 2012 (*1*). Recent exciting studies have begun to identify critical spinal cord inhibitory neurons and excitatory neurons in the transduction and modulation of mechanical itch signaling (*2–5*). Moreover, alloknesis can also be promoted by a loss of the inhibitory Piezo2-Merkel cell signaling as well as the inhibitory CD26-mediated dipeptidyl peptidase IV (DPPIV) enzyme activity in the skin (*6, 7*). However, the cellular and molecular mechanisms mediating mechanical itch in the skin are incompletely understood.

Recent RNA sequencing studies have revealed a subpopulation of TRPV1-expressing (TRPV1^+^) neurons also express Mas-related G-protein coupled receptor member A3 (MrgprA3) (*8*) and serve as a direct target for histamine and chloroquine (CQ), and mediates chemical-induced itch sensation in mice. Moreover, MrgprA3-TRP channel signaling triggers itch sensation but MrgprA3-P2X3 signaling produces pain sensation, indicating that MrgprA3 signaling functions as a polymodal signal integrator to allow the diversification of somatosensation (*9*). Interestingly, although *in vivo* electrophysiological recordings showed that MrgprA3^+^ neurons respond to mechanical stimuli (*10*), the molecular basis of the mechanosensitivity in the MrgprA3^+^ neurons remains unknown and the physiological/pathological relevance of MrgprA3^+^ neurons in mechanosensation also needs to be determined.

Piezo2 channel is a bona fide mechanosensitive ion channel and mediates the rapidly adapting (RA) mechanically activated currents in response to mechanical stimuli in various cell types (*10*– *14*). In addition to its expression in Merkel cell and thickly myelinated slowly-adapting type I (SAI) fibers, Piezo2 channel expressed by thinly myelinated Aδ and unmyelinated C afferents mediates mechanical hypersensitivity in response to tissue inflammation and peripheral nerve injury (*11, 15–17*). In contrast, whether Piezo2 channel is involved in chemical itch and/or mechanical itch is not well understood and the cellular basis of Piezo2 in the development of itch sensation remains unexplored.

Here we show that TRPV1^+^/MrgprA3^+^ neurons are required for mechanical itch induced by both histamine and CQ. Importantly, alloknesis-associated mechano-hypersensitivity in the TRPV1^+^/MrgprA3^+^ neurons relies on the sensitization of the Piezo2 channel, via Gαq/PLC/PKCδ signaling downstream of the activation of histamine receptors and MrgprA3, initiating mechanical itch but not acute chemical itch. Collectively, our data demonstrates that MrgprA3^+^ prurioceptor-Piezo2 signaling is essential to the generation of pruritogen-induced alloknesis.

## RESULTS

### Chemogenetic activation of TRPV1^+^ neurons promotes mechanical itch

Large-scale single-cell RNA sequencing revealed three distinct subtypes of nociceptive DRG neurons innervating the skin: the non-peptidergic polymodal C fibers expressing MrgprD, the peptidergic polymodal C fibers expressing TRPV1 and the low-threshold mechanosensitive C fibers (C-LTMRs) expressing tyrosine hydroxylase (TH) and vesicular glutamate transporter type 3 (Vglut3) (*18*). To determine the classes of DRG neurons involved in the generation of mechanical itch, we first investigated if chemogenetic stimulation of distinct sensory neuron subpopulations is sufficient to cause mechanical itch. We thus generated Trpv1-hM3Dq, Th-hM3Dq, and MrgprD-hM3Dq mice by crossing *Trpv1^Cre^*, *Th^CreER^* and *MrgprD^CreERT^* with *Rosa26^CAG-ds-hM3Dq^*mice, respectively. Intradermal injection of the DREADD agonist clozapine N-oxide (CNO) into the nape of neck induced a robust scratching response in the *Cre^+^* but not *Cre^-^ Trpv1-hM3Dq* mice (Supplementary Fig. 1). Interestingly, 30 min after the CNO injection when the spontaneous scratching subdued, a light touch stimulation delivered by the 0.7 mN von frey hair filament induced significantly increased mechanical itch in the *Cre^+^ Trpv1-hM3Dq* mice when compared with their *Cre^-^*littermates (Fig. 1A), recapitulating the alloknesis phenotype produced by intradermal injections of pruritogens (*1*). Surprisingly, after Cre induction by intraperitoneal injections of tamoxifen for 5 consecutive days, neither Th-hM3Dq nor MrgprD-hM3Dq mice showed increased mechanical itch in response to CNO treatment (Fig. 1B-1C). Collectively, these results demonstrate that TRPV1-lineage neurons but not mechanosensitive primary afferents expressing TH or MrgprD are critically involved in the initiation of mechanical itch.

**Figure 1.**
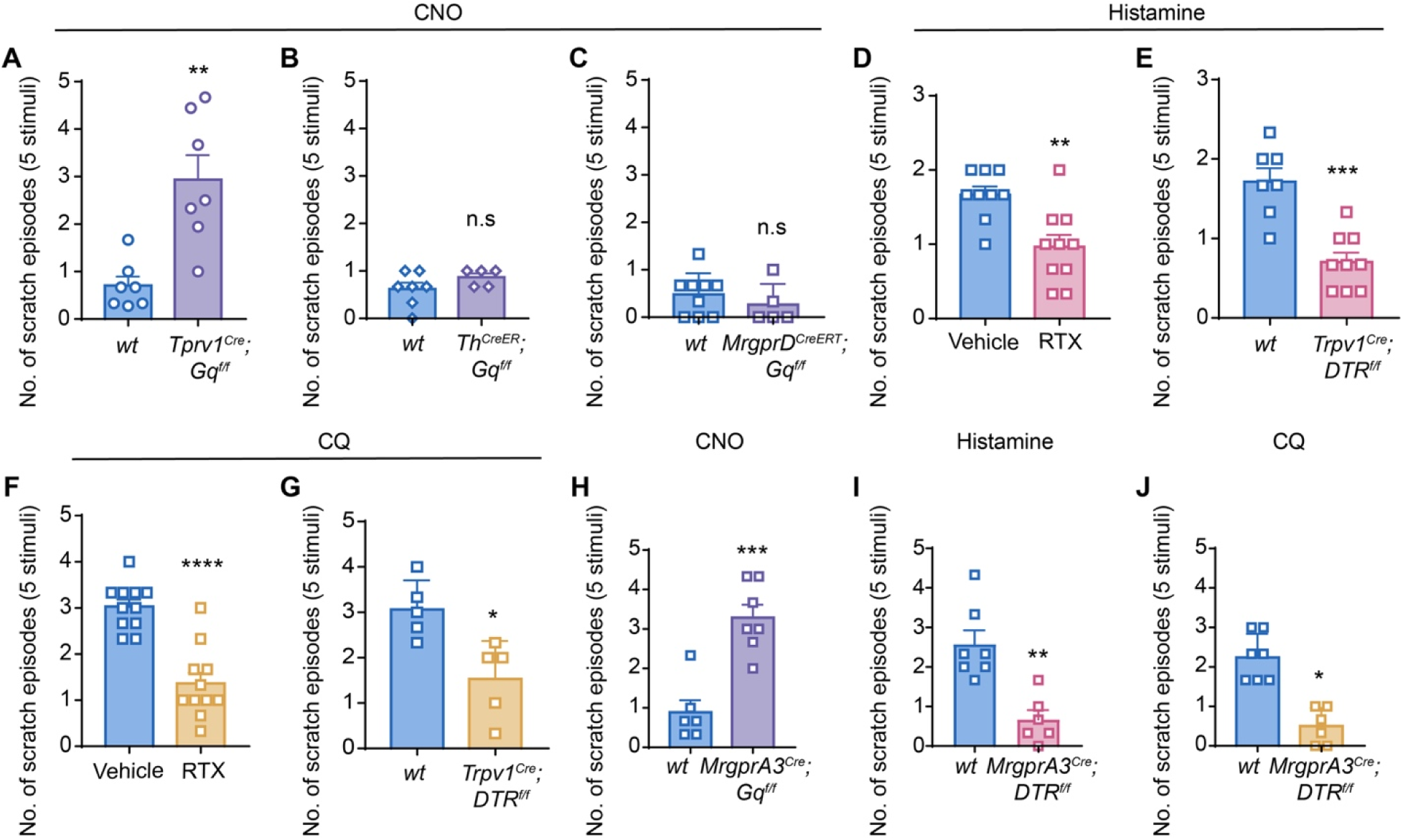
Trpv1^+^/MrgprA3^+^ neurons are critically involved in the initiation of mechanical itch. **A,** CNO-induced alloknesis score in the *Cre^-^* and *Cre^+^ Trpv1-hM3Dq* mice. n=7 for each group. **B**, CNO-induced alloknesis score in the *Cre^-^* (n=7) and *Cre^+^* (n=5) *Th-hM3Dq* mice. **C**, CNO-induced alloknesis score in the *Cre^-^* (n=9) and *Cre^+^* (n=5) *MrgprD-hM3Dq* mice. **D**, Histamine-induced alloknesis score in RTX-treated mice (n=10) and vehicle-treated littermates (n=9). **E**, Histamine-induced alloknesis score in the *Cre^-^* (n=7) and *Cre^+^* (n=9) *Trpv1-DTR* mice. **F**, CQ-induced alloknesis score in RTX-treated mice (n=11) and vehicle-treated littermates (n=11). **G**, CQ-induced alloknesis score in the *Cre^-^* (n=5) and *Cre^+^* (n=5) *Trpv1-DTR* mice. **H**, CNO-induced alloknesis score in the *Cre^-^* (n=6) and *Cre^+^*(n=7) *MrgprA3-hM3Dq* mice. **I**, Histamine-induced alloknesis score in the *Cre^-^* (n=7) and *Cre^+^* (n=6) *MrgprA3-DTR* mice. **J**, CQ-induced alloknesis score in the *Cre^-^* (n=7) and *Cre^+^* (n=6) *MrgprA3-DTR* mice. Data was shown as mean ± SEM. n.s, not significant. \**P*<0.05, \*\**P*<0.01, \*\*\**P*<0.001, \*\*\*\**P*<0.0001. Unpaired two-tailed Student’s t test.

### Chemically or genetically induced ablation of TRPV1^+^ neurons reduces histamine- and CQ-induced mechanical itch

To confirm the role of TRPV1^+^ neurons in the generation of pruritogen-induced alloknesis, we employed loss of function studies ablating TRPV1^+^ nociceptors by either intradermal injections of a super potent TRPV1 ligand resiniferatoxin (RTX) to wild type (*wt*) *C57bl/6* mice or intraperitoneal injections of diphtheria toxin (DTX) to the *TRPV1^Cre^*; *ROSA26^DTR^* mice. These ablation models were validated by a severely reduced thermal pain behavior in the tail-flick test (Supplementary Fig. 2). As expected, acute scratching behaviors evoked by either histamine or CQ were significantly reduced in RTX-treated mice and DTX-treated *Cre^+^ TRPV1^Cre^*; *ROSA26^DTR^* mice when compared with their respective control mice, indicating an efficient ablation of TRPV1^+^ neurons (Supplementary Fig. 3A-3D).

30 min after the spontaneous scratching induced by histamine or CQ subdued, we delivered light touch stimuli and analyzed the alloknesis scores in histamine- and CQ-injected mice. Surprisingly, in contrast to a previous study showing that CQ did not evoke alloknesis at the concentration of 200 nmol/10 μl (*1*), intradermal injections of CQ evoked mechanical itch in a bell-shaped manner with the peak effect produced by the concentration of 50 nmol/30 μl (Supplementary Fig. 4A-4D), suggesting a higher concentrations of CQ may lead to an inhibition and/or desensitization of alloknesis. Both histamine-induced and CQ-induced mechanical itch was markedly reduced in RTX-treated and DTX-treated *Cre^+^ TRPV1^Cre^*; *ROSA26^DTR^*mice when compared with their control littermates (Fig. 1D-1G). Moreover, silencing the TRPV1^+^ neurons by selective delivery of a charged membrane-impermeable sodium channel blocker lidocaine N-ethyl-lidocaine (QX-314) plus CQ also significantly reduced CQ- and histamine-induced mechanical itch (Supplementary Fig. 5A and 5B). Taken together, these results suggest that TRPV1^+^ C-fibers are necessary and sufficient to produce pruritogen-induced mechanical itch.

### MrgprA3^+^ pruriceptors mediate mechanical itch

A subpopulation of TRPV1^+^ and histamine H1 receptor^+^ neurons express the CQ receptor MrgprA3 and have been well-characterized to play important roles in both acute and chronic itch (*8, 19, 20*). However, whether MrgprA3^+^ prurioceptors are involved in the pathogenesis of pruritogen-induced alloknesis remains unknown. To address this question, we expressed the excitatory Gq-DREADD in the MrgprA3^+^ neurons by crossing *MrgprA3^GFP-Cre^* mice with *Rosa26^CAG-ds-hM3Dq^*. Intradermal injections of CNO in the *Cre^+^ MrgprA3-hM3Dq* mice produced a robust spontaneous scratching behavior while their *Cre^-^* littermates didn’t show any itch response (Supplementary Fig. 6A), supporting functional expression of hM3Dq in the MrgprA3^+^ prurioceptors. Interestingly, light mechanical stimulation evoked mechanical itch responses were greatly increased in the *Cre^+^* but not in the *Cre^-^ MrgprA3-hM3Dq* mice 30 minutes after CNO injection (Fig. 1H), recapitulating the mechanical itch response induced by intradermal injections of CQ. Corroborating with the gain of function experiments, selectively ablation of the MrgprA3^+^ neurons by DTX in the *MrgprA3^cre^; ROSA26^DTR^* mice greatly attenuated histamine- and CQ-induced acute chemical itch (Supplementary Fig. 6B-6C) and mechanical itch (Fig. 1I-1J) in the *Cre^+^MrgprA3^cre^; ROSA26^DTR^* mice when compared with that in their *Cre^-^* littermates. Taken together, our results suggest that MrgprA3^+^ neurons are sensitized by histamine and CQ, contributing to the development of mechanical itch in addition to acute chemical itch.

### Piezo2 channel but not TRPA1 channel is required for pruritogen-induced mechanical itch

Although it is commonly accepted that TRPA1 is expressed by a subset of TRPV1^+^ nociceptors and acts as a chemosensor mediating both inflammatory and neuropathic pain as well as pruritogen-induced non-evoked spontaneous scratching, the role of TRPA1 in mechanosensation has been controversial (*21–24*). We therefore investigated whether TRPA1 is also involved in pruritogen-induced mechanical itch. Surprisingly, comparable alloknesis scores were found between the *TRPA1^+/+^* and *TRPA1^-/-^* mice in the presence of either histamine or CQ (Supplementary Fig. 7), suggesting that TRPA1 is not involved in the development of pruritogen-induced alloknesis.

On the other hand, although we have demonstrated that epithelia Merkel cell-expressing Piezo2 channel modulates mechanical itch through driving the Merkel cell-SA1 Aβ signaling (*6*), the role of mechanosensitive Piezo2 channel expressed by the MrgprA3^+^ C-type pruriceptors is not known. To address this, we first determined the expression of Piezo2 in the MrgprA3^+^ neurons using Piezo2 mRNA fluorescent *in situ* hybridization in DRG from the *MrgprA3^GFP^*^-*Cre*^ mice. Indeed, 12.5% of Piezo2-expressing neurons were labeled by MrgrpA3-GFP while more than 80% of MrgprA3-GFP^+^ neurons expressed Piezo2 (Fig. 2A and 2B). To determine if Piezo2 is functionally expressed by MrgprA3^+^ neurons, we crossed the *MrgprA3^GFP-cre^; Ai9^f/f^* mice with *Piezo2^f/f^* mice and performed whole-cell patch-clamp recording. Mechanically-activated (MA) whole-cell inward currents were recorded in 10 out of 32 DRG neurons from 3 *Cre^-^MrgprA3^cre^; Ai9^f/f^* mice while none of 28 neurons from 3 *Cre^+^ MrgprA3^cre^; Ai9^f/f^; Piezo2^f/f^* mice responded to the same stimuli (Fig. 3C and 3D), suggesting Piezo2 channel confers mechanosensitivity to MrgprA3^+^ pruriceptors.

**Figure 2.**
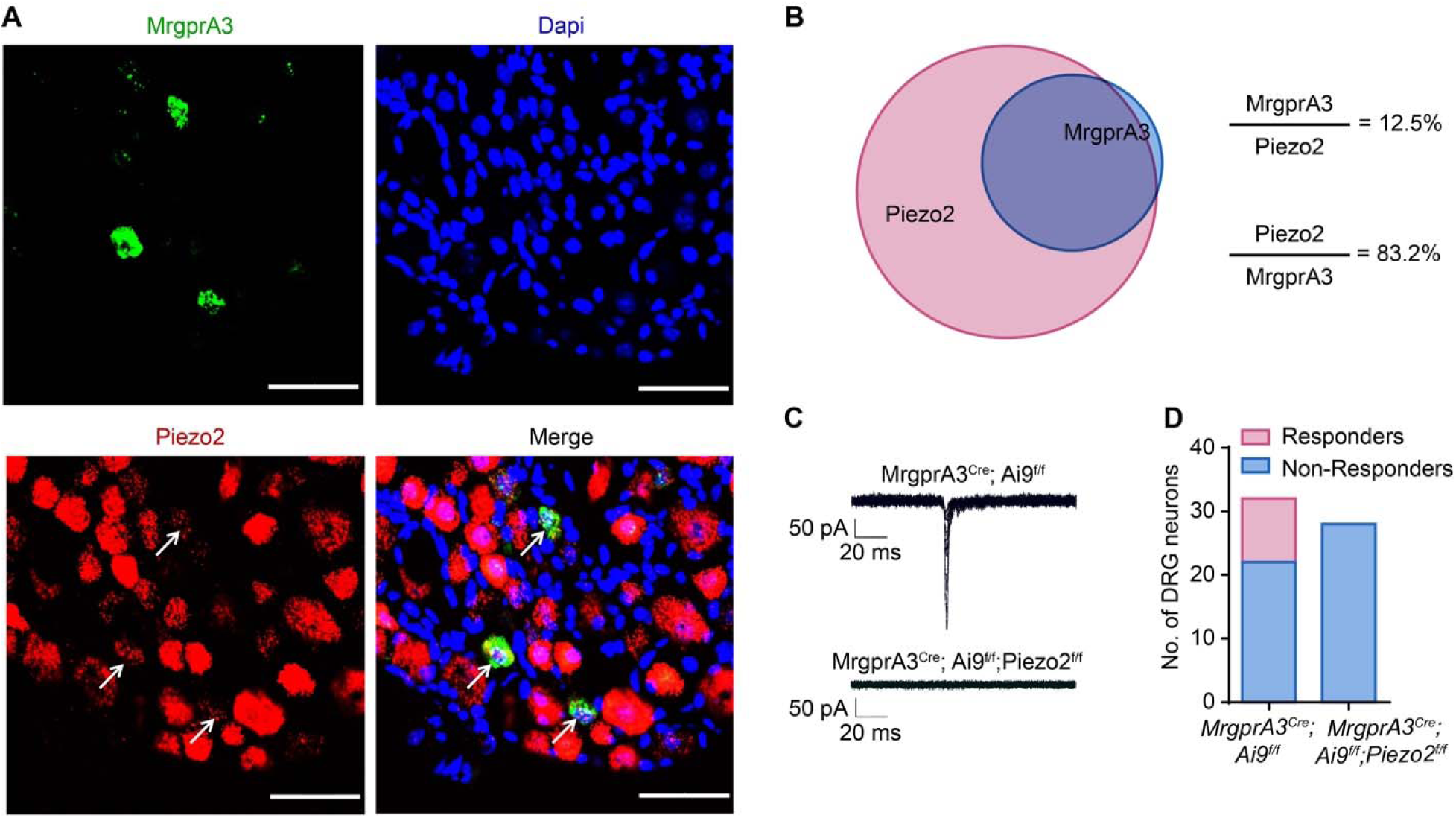
Characterization of Piezo2 expression in the MrgprA3^+^ neuron. **A,** *In situ* hybridization staining of MrgprA3-GFP signals (green) with Piezo2 (red). Arrows indicate double-labeled cells. n=3-5 sections from 3 mice. Scale bar, 50 µm. **B,** The BioVenn diagram illustrates the overlap between MrgrprA3^+^ neurons and Piezo2^+^ neurons. **C,** Representative whole-cell MA current traces elicited by mechanical indentation on DRG neurons isolated from *MrgprA3^cre^; Ai9^f/f^* and *MrgprA3^cre^; Ai9^f/f^; Piezo2^f/f^* mice. **D,** Bar chart summarized the mechanosensitivity of recorded DRG neurons isolated from *MrgprA3^cre^; Ai9^f/f^* and *MrgprA3^cre^; Ai9^f/f^; Piezo2^f/f^*mice.

**Figure 3.**
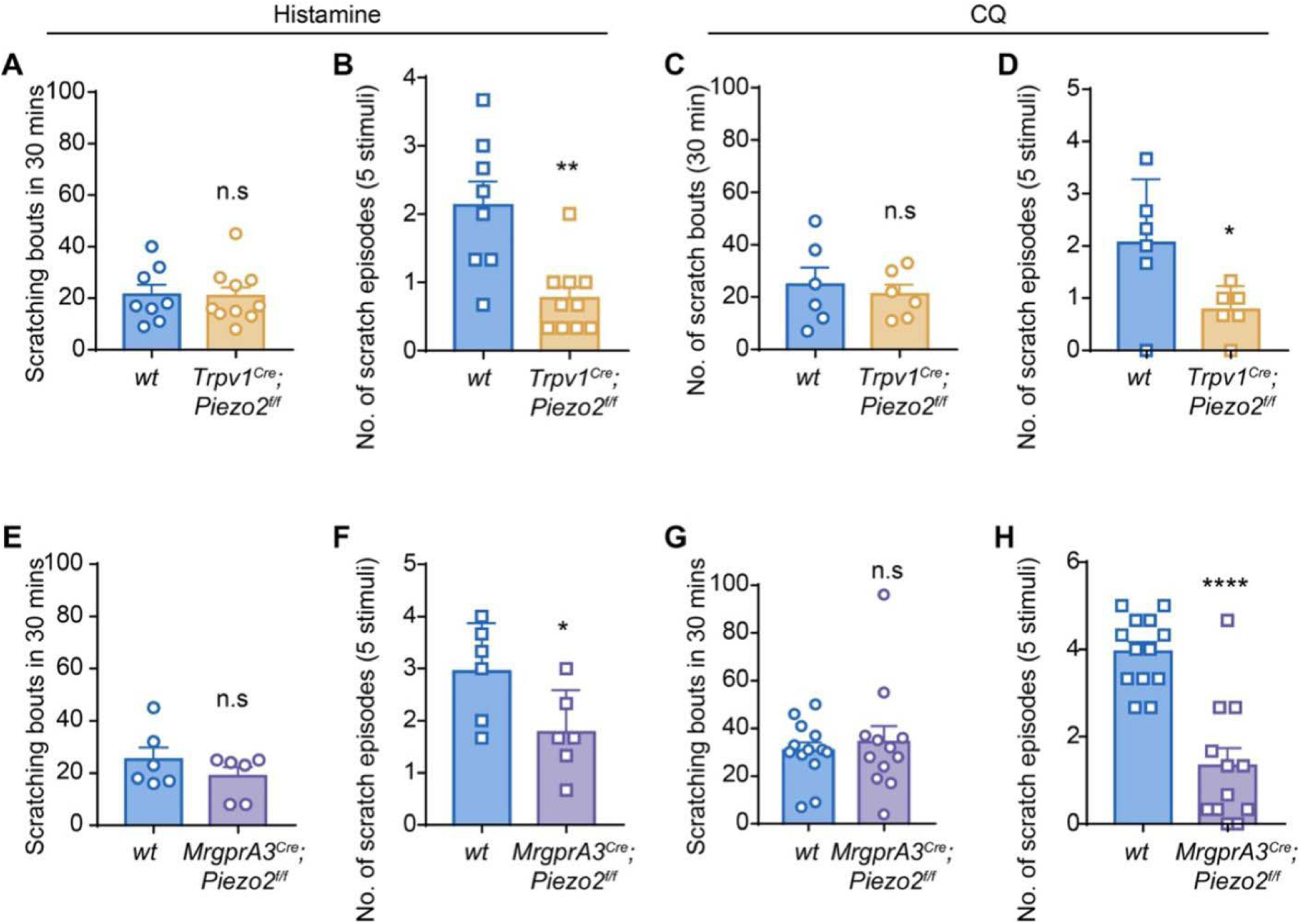
Pruriceptor-expressing Piezo2 is required for histamine and CQ induced mechanical itch. **A-B** Intradermal injection of histamine induced acute itch (A) and mechanical itch (B) in the *Cre^-^* (n=8) and *Cre^+^*(n=10) *Trpv1^Cre^*; *Piezo2^f/f^* mice. **C-D** Intradermal injection of CQ induced acute itch (C) and mechanical itch (D) in the *Cre^-^*(n=6) and *Cre^+^* (n=6) *Trpv1^Cre^*; *Piezo2^f/f^* mice. **E-F** Intradermal injection of histamine induced acute itch (E) and mechanical itch (F) in the *Cre^-^*(n=6) and *Cre^+^* (n=6) *MrgprA3^Cre^*; *Piezo2^f/f^* mice. **G-H** Intradermal injection of CQ induced acute itch (G) and mechanical itch (H) in the *Cre^-^* (n=13) and *Cre^+^* (n=12) *MrgprA3^Cre^*; *Piezo2^f/f^* mice. Data was shown as mean ± SEM. n.s, not significant. \**P*<0.05, \*\**P*<0.01, \*\*\*\**P*<0.0001. Unpaired two-tailed Student’s t test.

To further investigate whether TRPV1^+^/MrgprA3^+^ pruriceptor-expressed Piezo2 channel is involved in the generation of pruritogen-induced mechanical itch, we injected CQ and histamine into the nape of the neck of the *TRPV1^Cre^*; *Piezo2^f/f^* mice and *MrgprA3^cre^; Piezo2^f/f^*mice. Although the acute chemical itch induced by either CQ or histamine was comparable between the *Cre^+^*and *Cre^-^ TRPV1^Cre^*; *Piezo2^f/f^* mice or *MrgprA3^cre^; Piezo2^f/f^* mice (Fig. 3A, 3C, 3E and 3G), alloknesis scores were markedly reduced in the C*re^+^* mice when compared with that in their respective *Cre^-^* littermates (Fig. 3B, 3D, 3F and 3H), suggesting that Piezo2 channel functions as a critical downstream mediator of MrgprA3 signaling in the generation of mechanical itch but not acute itch produced by pruritogens.

### PLC-PKC**δ** signaling sensitizes Piezo2 channel to mediate pruritogen-induced mechanical itch

Prior studies demonstrated that increased levels of inflammatory mediators in the inflamed or injured tissues may sensitize Piezo2 function, leading to enhanced mechanical sensitivity (*11, 15, 17*), which involves multiple intracellular signaling pathways (*16, 25, 26*). Corroborating with these findings, both applications of CQ and histamine significantly increased the MA current density in DRG neurons from *Cre^+^ MrgprA3^cre^; Ai9^f/f^* mice (Fig. 4A and 4B), suggesting that pruritogen applications sensitize Piezo2 channel function in the MrgprA3^+^ neurons.

**Figure 4.**
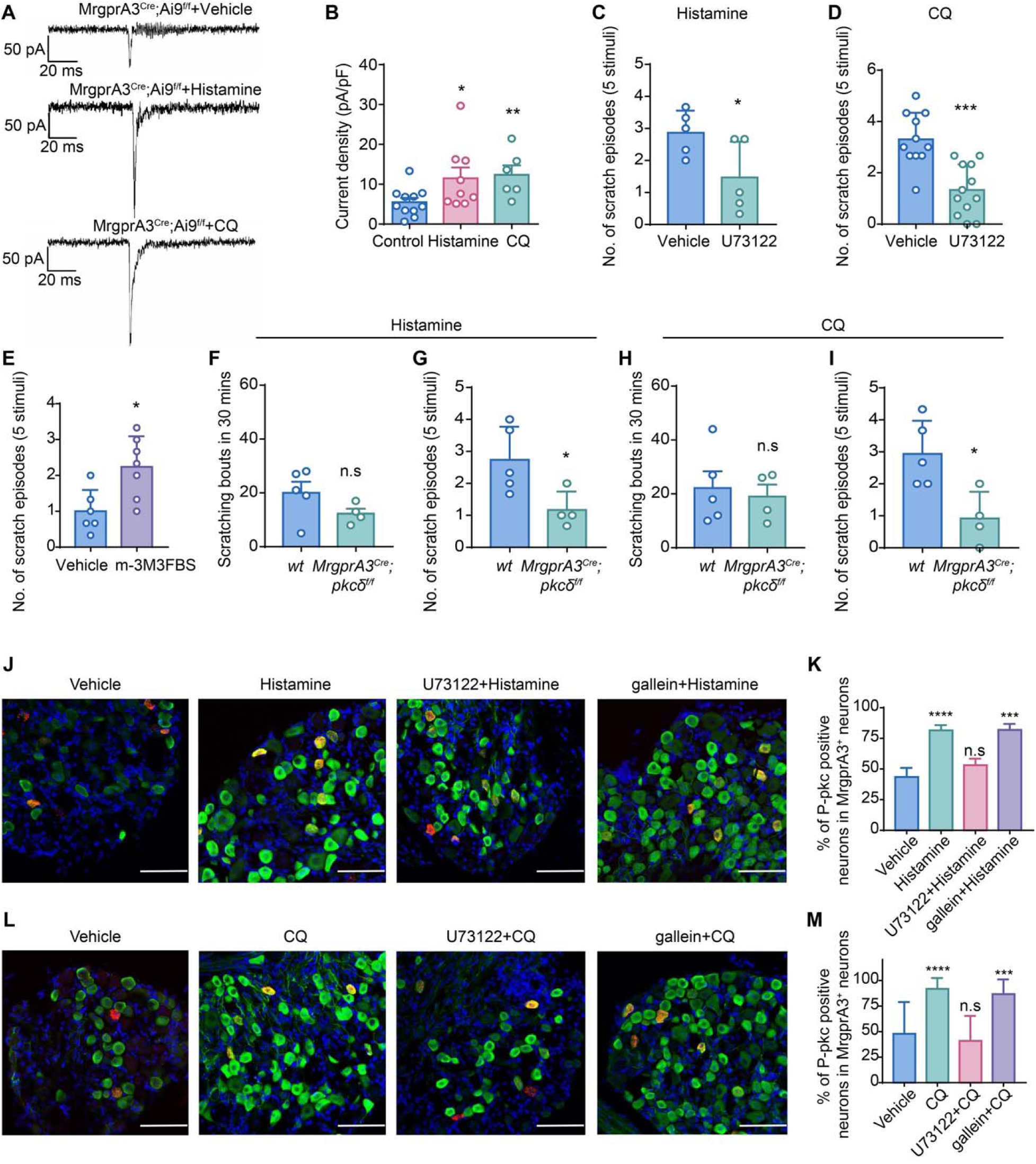
Pruritogens sensitize Piezo2 through PLC-PKCδ signaling in mechanical itch. **A,** Representative whole-cell MA current traces elicited by mechanical indentation on DRG neurons isolated from *MrgprA3^cre^; Ai9^f/f^* mice perfused with vehicle, histamine or CQ, respectively. **B,** Summarized data of the current density of mechanosensitive MrgprA3^+^ neurons in response to vehicle (n=11), histamine (n=9) and CQ (n=6). **C-D** Histamine-induced (C) and CQ-induced (D) alloknesis score in mice treated with vehicle or the PLC inhibitor U73122. n=5 mice for each group in C, n=11 vehicle-treated and n=12 U73122-treated mice in D. **E**, The selective PLC agonist induced alloknesis score in *wt* mice. n=6 vehicle-treated and n=7 m-3M3FBS-treated mice. **F-G**, Intradermal injection of histamine-induced acute itch (F) and mechanical itch (G) in the *Cre^-^*(n=5) and *Cre^+^* (n=4) *MrgprA3^Cre^*; *PKC*δ*^f/f^* mice. **H-I**, Intradermal injection of CQ- induced acute itch (H) and mechanical itch (I) in the *Cre^-^* (n=5) and *Cre^+^*(n=4) *MrgprA3^Cre^*; *PKC*δ*^f/f^* mice. **J**, Representative immunofluorescent staining of p-PKC in DRG neurons isolated from *MrgprA3^Cre-GFP^* mice treated with vehicle, Histamine, U73122 + Histamine, gallein + Histamine. **K,** Quantification of number of p-PKC expressing cells in MrgprA3^+^ neurons in the present of different treatments in J, n=3-5 sections from 3 mice for each group. **L**, Representative immunofluorescent staining of p-PKC in DRG neurons isolated from *MrgprA3^Cre-GFP^* mice treated with vehicle, CQ, U73122 + CQ, gallein + CQ. **M**, Quantification of number of p-PKC expressing cells in MrgprA3^+^ neurons in the present of different treatments in L. n=3-5 sections from 3 mice for each group. Green, p-PKC; red, MrgprA3. All scale bar, 100 µm. Data was shown as mean ± SEM. n.s, not significant. \**P*<0.05, \*\**P*<0.01, \*\*\**P*<0.001, \*\*\*\**P*<0.0001. Unpaired two-tailed Student’s t test in C-I. One-way ANOVA in K and M.

GPCRs including MrgprA3 and histamine receptors could be dissociated into G_αq/11_ and G_βγ_ subunits, mediating noxious sensation through coupling to downstream ion channels. To determine the signaling pathway involved in Piezo2 channel sensitization, we first employed gallein, a small molecule inhibitor of G_βγ_ which has been shown to mediate TRPA1-dependent itch (*27, 28*). Surprisingly, pretreatment of gallein blocked neither the chemical itch nor the mechanical itch induced by CQ or histamine (Supplementary Fig. 8).

On the other hand, it was also reported that CQ-induced scratching behavior and neuronal activities were abolished in the Phospholipase C beta 3 (PLCβ3) knockout mice (*29*). To test whether the PLC signaling contributes to MrpgrA3^+^ neuron sensitization in mechanical itch, we pre-treated *wt* mice with the PLC inhibitor U73122 and PLC agonist m-3M3FBS. Surprisingly, U73122 significantly reduced histamine- and CQ-evoked mechanical itch (Fig. 4C and 4D) while application of m-3M3FBS alone was sufficient to promote mechanical itch (Fig. 4E), suggesting that PLC signaling is necessary and sufficient to the generation of mechanical itch.

Upon its activation, PLC hydrolyzes phosphatidylinositol 4,5-bisphosphate (PIP2) into inositol 1,4,5-trisphosphate (IP3) and 1,2-diacylglycerol (DAG). While IP3 is critical to intracellular Ca^2+^ mobilization, the main effect of DAG is to activate PKC, a family of serine/threonine kinases comprising 11 isoforms encoded by 9 genes (*30*). To determine the PKC isoform involved in MrgprA3-dependent alloknesis, we first re-analyzed the single-cell RNA-seq data of DRG cells (*20*) and found that PKCδ is the highest expressed PKC isoform in MrgprA3^+^ neurons (supplementary Fig 9). Thus, we genetically ablated PKCδ from the MrgprA3^+^ neurons by crossing the *MrgprA3^cre^* mice with *PKC*δ*^f/f^* mice. Interestingly, although pruritogen-induced acute chemical itch was comparable, mechanical itch evoked by CQ or histamine was significantly reduced in the *PKC*δ*^cko^* mice when compared with that in the littermate controls (Fig. 4F-4I), suggesting that activation of PKCδ downstream of MrgprA3 and histamine receptor signaling sensitizes the mechanosensitivity in the MrgprA3^+^ neurons. Thus, an intracellular PLC- PKCδ-Piezo2 signaling contributes to pruritogen-induced mechanical itch sensitization.

Moreover, previous studies also revealed that priming by phosphorylation, PKC is activated by PLC signaling (*31, 32*) and phospho-PKC (p-PKC) represents a sensitive biomarker of neuronal activity (*33*). Consistent with this note, our immunofluorescent staining revealed an increased p-PKC expression in response to histamine- or CQ-induced activation of the MrgprA3^+^ neurons, which was blocked by U73122 but not gallein (Fig. 4J-4M), further indicating that PKC activation is tightly correlated with pruritogen-induced MrgprA3^+^ neuron excitability.

## DISCUSSION

Mechanical itch (alloknesis) is an exaggerated itch sensation elicited by innocuous mechanical stimuli, especially in the presence of exogenous pruritogens or in the setting of chronic itch. Although alloknesis was first described 30 years ago (*34, 35*), the molecular and cellular mechanisms underlying pruritogen-induced alloknesis are not well understood. Here we provided multiple lines of evidence showing that TRPV1^+^/MrgprA3^+^ pruriceptors mediate mechanical itch following administration of pruritogens CQ and histamine, which requires the function of mechanosensitive Piezo2 channel. We further demonstrated that GPCR-PLC-PKCδ signaling induced by CQ or histamine sensitizes Piezo2 channel function to produce mechanical itch (Supplementary Fig. 10). Our results place the TRPV1^+^/MrgprA3^+^ pruriceptors-Piezo2 signaling axis at the center of mechanical itch signaling pathway driven by exogenously applied pruritogens.

Published studies have shown multiple forms of mechanical itch under both physiological and pathological conditions. For instance, Pan et al. identified a subpopulation of Ucn3^+^ excitatory interneurons in the spinal cord which are required for mechanical itch under physiological conditions(*9*). In the absence of pruritogens or chronic itch conditions, Ucn3-expressing excitatory interneurons as well as the neuropeptide Y1 receptor-expressing excitatory interneurons in the spinal cord are required for the transmission of mechanical itch occurring in specific locations (*2*), i.e., skin areas behind the ears. Interestingly, these Ucn3^+^ neurons were shown to receive TLR5^+^ Aβ LTMRs which express NF200 and contribute to the generation of touch and mechanical allodynia (*5, 36*). Besides potential role of the TLR5-Ucn3 signaling in mediating mechanical itch under physiological conditions, mechanical itch can also be evoked in the setting of chronic skin inflammation such as that in mouse models of experimental dry skin (*1*) and imiquimod-induced psoriasis (*37*) as well as exogenously applied pruritogens such as histamine (*38*) and endomorphins (*7*).

In addition to TLR5^+^ Aβ LTMRs, several C-LTMRs have been shown to play important roles in mechanical hypersensitivity including C fibers expressing Vglut3/TH and MrgprD. MrgprD^+^ DRG neurons constitute >90% of all nonpeptidergic cutaneous C-fibers but are not overlapping with the TRPV1^+^ nociceptors (*39*). Although activation of MrgprD^+^ neurons evoke acute itch in mice and MrgprD^+^ neurons exhibit enhanced excitability in a mouse model of chronic itch associated with allergic contact dermatitis (ACD) (*40–42*), our results showed that chemogenetic activation of MrgprD^+^ neurons does not evoke mechanical itch, suggesting that MrgprD^+^ nonpeptidergic C polymodal nociceptors are unlikely involved in pruritogen-induced mechanical itch in mice. In addition to MrgprD^+^ C polymodal nociceptors, Vglut3^+^/TH^+^ C-LTMRs are also not overlapping with the TRPV1^+^ peptidergic cutaneous C-fibers, mediating mechanical allodynia in the setting of neuropathic pain (*43, 44*). On the other hand, our genetic studies found no significant effect of the Vglut3^+^/TH^+^ C-LTMRs on mechanical itch. Interestingly, a recent study demonstrated that the Vglut3-lineage neurons mediate spinal inhibition of mechanical itch (*45*), suggesting that activation of the Vglut3^+^/TH^+^ C-LTMRs does not contribute to pruritogen-induced mechanical itch but inhibits mechanical itch in the spinal circuit instead. Despite these findings exclude the involvement of the MrgprD^+^ and Vglut3^+^/TH^+^ mechanosensitive receptors in pruritogen-induced mechanical itch, our results provide conclusive evidence that the TRPV1^+^/MrgprA3^+^ C pruriceptors are necessary and sufficient for pruritogen-induced mechanical itch as demonstrated in both gain- and loss-of-function studies.

Chemically induced acute itch (spontaneous itch) elicits the scratching reflex in a non-evoked manner while mechanical-induced itch requires mechanical stimuli to initiate the subsequent scratching behavior. Furthermore, chemically-induced itch declined over a 30-min period while mechanically-induced itch lasted for at least 2 hours. Thus, the differences between chemical itch and mechanical itch may implies distinct signaling pathway. Indeed, although TRPA1 has been shown to act downstream of the CQ-MrgprA3 signaling, global TRPA1 knockout mice did not show differences in either histamine- or CQ-induced mechanical itch, which is consistent with previous studies (*46, 47*). In marked contrast, our study demonstrated that ablation of Piezo2 channel function from the MrgprA3^+^ neurons diminishes mechanical itch induced by both CQ and histamine, suggesting a critical role of MrgprA3-Piezo2 signaling in mechanical itch associated with pruritogen-mediated sensitization of pruriceptive neurons. These results also suggest that the intrinsic mechanosensitivity of MrgprA3^+^ neurons contributing to the generation of mechanical itch is likely mediated by Piezo2 channel.

It should be noted that GPCR downstream signaling pathways couple various ion channels and contribute to distinct pruritogen-induced itch sensation. For instance, prior studies showed that histamine elicits acute itch sensation by activating downstream TRPV1 through PLCβ3 and PLA2/lipoxygenase (*29, 48, 49*); CQ evokes TRPA1-dependent itch through a signaling mechanism involving MrgprA3-coupled Gβγ signaling but not PLC (*50*). Moreover, it was reported that inhibition of PKCδ significantly decrease histamine-induced acute itch, mainly because PKCδ is phosphorylated in response to histamine treatment and contributes to the histamine-mediated sensitization of the voltage-gated sodium channel Nav1.7 (*51, 52*). Interestingly, our results further showed that PLC-PKCδ signaling-mediated sensitization of MrgprA3^+^ neuron-expressing Piezo2 channel play pivotal role in pruritogen-induced mechanical itch but not acute chemical itch, revealing a novel signaling axis of PLC-PKCδ-Piezo2 in mechanical itch, which is also distinct from the EPAC1-Piezo2 (*26*) and PKA/PKC-Piezo2 signaling axis in mechanical pain (*15*) .

In summary, our findings show that activation of TRPV1^+^/MrgprA3^+^ DRG neurons mediates pruritogen-induced mechanical itch by sensitizing mechanosensitive Piezo2 channel through PLC signaling in the skin. Identification of the MrgprA3^+^ C pruriceptors in mediating the pruritogen-induced mechanical itch highlights the importance of this small subset of TRPV1^+^ neurons in the generation of both mechanically stimulated and non-evoked spontaneous itch not only to the MrgprA3 receptor ligand CQ but also to other pruritogens such as histamine. Our data shed light on the TRPV1/MrgprA3-PLC-PKCδ-Piezo2 signaling axis in the development of alloknesis which may offer a novel therapeutic target for treating chronic itch.

## ACKNOWLEDGMENTS

We thank Dr. Mark Hoon for sharing the *TRPV1^Cre^* mice and Dr. Xinzhong Dong for sharing the *MrgprA3^GFP-Cre^*mouse line. This work is supported by grant from National Institutes of Health grant R01AA027065, R01AR077183 and R01DK103901 to H.Z.H and R01NS106289 to G.F.W.

## AUTHOR CONTRIBUTIONS

H.H and J.F conceived of the project and wrote the manuscript. P.L designed and conducted most of mouse behavioral experiments. P.L and Y.Z was responsible for RNAscope and immunostaining. Y.Z and Z.X was responsible for patch clamp recordings. Others assisted with the data analysis and manuscript preparation.

## DATA AND CODE AVAILABILITY

This study did not generate/analyze datasets or code.

## MATERIALS AVAILABILITY

This study did not generate new unique reagents. Further information and requests for resources and reagents should be directed to and will be fulfilled by Dr. Hongzhen Hu and Dr. Jing Feng.

## STAR METHODS

### Mice

All animal procedures were performed using protocols approved by the Animal Studies Committee at Washington University School of Medicine and in compliance with the guidelines provided by the National Institute of Health and the International Association for the Study of Pain. Both adult male and female mice (8–12 weeks old) were used for all experiments and mice were randomly assigned to different experimental conditions. Mice were sex- and age-matched in all experiments and were group-housed in standard mouse housing cages at room temperature with unrestricted access to food and water on a 12 h light/12 h dark cycle.

We purchased *C57BL/6J* (Strain #: 000664)*, Th^CreER^* (Strain #: 008532)*, MrgprD^CreERT^*(Strain #: 031286), *Gq-DREADD* (*B6N;129-Tg^(CAG-CHRM3, mCitrine)1Ute/J^,* Strain #: 02622*0*), *ROSA26^iDTR^* (Strain #: 007900), Ai9 (*B6;129S6-Gt (ROSA)26Sor^tm9(CAG-tdTomato)Hze/^J*, Strain #: 007909), *Piezo2^flox/flox^*(Strain #: 027720) and *Trpa1^−/−^* (Strain #: 006401) mouse lines from Jackson Laboratory. *PKC*δ*^flox/flox^* (Strain #: 06462) mice were obtained from RIKEN BRC. *TRPV1^Cre^*mice were donated by Dr. Mark Hoon from NIH. The transgenic *MrgprA3^GFP-Cre^* mouse line was kindly provided by Xinzhong Dong (Johns Hopkins University, HHMI). For lineage tracing, Ai9 (*B6;129S6-Gt (ROSA)26Sor^tm9(CAG-tdTomato)Hze/J^*) were crossed with *MrgprA3^GFP-Cre^* to induce tdTomato expression in MrgprA3^+^ cells. *Piezo2^flox/flox^* mice were mated with *Trpv1^Cre^* and *MrgprA3^GFP-Cre^* mice to generate Piezo2 conditional knockout mice (*Piezo2^cKO^*). *PKC*δ*^flox/flox^* mice were mated with *MrgprA3^GFP-Cre^* mice to generate PKCδ conditional knockout mice (*PKC*δ*^cKO^*). The *Rosa26^iDTR^* mice were crossed with *Trpv1^Cre^* and *MrgprA3^GFP-Cre^*mice to obtain *Trpv1^Cre^*; *iDTR* and *MrgprA3^GFP-Cre^*; *iDTR* mice. For chemogenetic activation of DRG neurons, the transgenic mice were engineered by crossing *Trpv1^Cre^, Th^CreER^, MrgprD^CreERT^* and *MrgprA3^GFP-Cre^* mice with *Gq-DREADD (B6N;129-Tg ^(CAG-CHRM3, mCitrine)1Ute/J)^)* mice, respectively.

### Pruritogens-induced acute itch

To test the acute itch behavior, the fur on the nape of the neck was shaved and mice were acclimated in the red transparent recording chamber for 5 days. On the testing day, mice were habituated in the behavioral testing apparatus for 1 hour and then histamine (50 μg; Sigma-Aldrich, St. Louis MO) or CQ (50 nmol; Sigma-Aldrich, St. Louis, MO) in 30 µL sterile saline was injected intradermally to the nape of the neck. Insulin syringes with 30G needle (UltiCare) were used for intradermal injection. Immediately after the injection, mice were videotaped for 30 min. After the recording, the videotapes were played back and the number of scratch bouts were counted over the 30-min recording period by an investigator blinded to the treatment. A scratching bout is defined as one or more rapid back-and-forth motion of the hindpaw directed to the site of injection, and ending with licking or biting of the toes and/or placement of the hindpaw on the floor.

### Pruritogens-induced alloknesis

Mice were acclimated in a red recording chamber with a removable mesh cover for at least 5 days. 30 min after intradermal injection of respective pruritogens, mice received an innocuous mechanical stimulus for 1 second delivered using a von Frey filament (bending force: 0.7 mN) at five randomly-selected points oriented radially 7 mm away from the injection site. Mice received 3 separate stimulations at each point with an interval of 10 seconds. The scratching response of hind paw toward the poking site was considered as a positive response. The scratching number at each point was averaged and then summed into the final alloknesis score for comparison.

### Drug administration

To induce robust Cre activity, tamoxifen (Sigma, St. Louis, MO, USA) was dissolved in corn oil and made fresh daily before use. Both *Cre^-^* and *Cre^+^*mice received intraperitoneal injection of tamoxifen at 100 mg/kg body weight for 5 consecutive days. *In vivo* experiments were performed between 7 and 14 days after tamoxifen injection.

For the chemogenetic stimulation, *Cre^+^* and *Cre^-^ mice* of *Trpv1; Gq-DREADD, Th; Gq-DREADD, MrgprD; Gq-DREADD* and *MrgprA3; Gq-DREADD* mice were intradermally injected with 50 ul 3 mM clozapine N-oxide CNO (Sigma-Aldrich, St. Louis, MO). Immediately after the injection, mice were videotaped for 30 min. Alloknesis scores were evaluated 0.5 hours after CNO injections.

### Pharmacological silence of sensory fibers in the skin

Intradermal injection of 30 μl 0.2% QX-314+0.3 µg flagellin or 30 μl 0.2% QX-314+ 50 nmol CQ are employed to block the A beta fibers or the C fibers, respectively. Flagellin (Sigma-Aldrich, St. Louis, MO), QX-314 (Sigma-Aldrich, St. Louis, MO) and CQ (Sigma-Aldrich, St. Louis, MO) were dissolved in isotonic saline. QX-314 (0.2%, 30 µl) was intradermally injected into the nape of the neck as a vehicle control. The effect of the Phospholipase C (PLC) activator m-3M3FBS (20 μM, 30 μl; Sigma-Aldrich, MO), PLC antagonist U73122 (0.1 μM, 30 μl; Sigma-Aldrich, MO) and the G protein βγ subunit inhibitor gallein (500 μM, 30 μl; Tocris Bioscience, USA) and on scratching and alloknesis was tested.

### In Situ Hybridization and Immunohistochemistry

Non-isotopic in situ hybridization (ISH) on DRG sections prepared from *MrgprA3^GFP-Cre^* mice was performed to detect Piezo2 mRNA expression. RNAscope Multiplex Fluorescent Reagent Kit V2 (Cat # 323100) and Piezo2 probes (400191) were purchased from ACDBio. The ISH/GFP double staining was performed as previously described (Liu et al., 2009). Briefly, after in situ hybridization for Piezo2 as instructed in the manual, DRG sections were incubated overnight at 4°C with chicken anti-GFP antibody (GFP-1020, Aves Labs, 1:1,000) and then incubated for 1 h at room temperature with donkey anti-chicken IgY (703-545-155, Alexa Fluor 488 conjugated, Jackson ImmuneResearch). Sections were then washed three times in PBS and mounted with DAPI (E140588, Invitrogen) for imaging.

To detect pruritogens-induced p-PKC expression in the MrgprA3^+^ neurons, *wt* mice were divided into eight groups for double staining. Four groups of mice were intradermally injected with vehicle (isotonic saline), CQ (50 nmol, 30 μl), U73122 (0.1 μM, 30 μl; 30 min prior to the CQ) + CQ and gallein (500 μM, 30 μl; 30 min prior to the CQ) + CQ respectively, and the other four groups were administered intradermally with vehicle (isotonic saline), histamine (50 µg, 30 μl), U73122 (0.1 μM, 30 μl; 30 min prior to the histamine) + histamine and gallein (500 μM, 30 μl; 30 min prior to the histamine) + histamine respectively. Thirty minutes after treatments, mice were deeply anesthetized with sodium pentobarbital (65 mg/kg, i.p.) and perfused through the heart with phosphate buffered saline (PBS, 0.1 M, pH 7.4) followed with fixative (4% paraformaldehyde, pH 7.4) at room temperature. Thoracic DRG were dissociated and fixed in fixative at 4 °C for overnight, then stored in 30% sucrose in PBS at 4 °C for 3 days. Samples were then embedded in optimal cutting temperature (OCT) compound (Sakura Finetek, Tissue-Tek, PA). Sections of approximately 12 μm on slides were performed for ISH with MrgprA3 probes (548161, ACDBio) firstly, then DRG sections were incubated overnight at 4°C with rabbit antibody to p-PKC (ab109539, Abcam, 1:500) and then incubated for 1 h at room temperature with goat anti-rabbit IgG H&L (ab150077, Alexa Fluor® 488, Abcam,).

Both fluorescent and ISH signals were observed under the Nikon C2 Confocal Laser Microscope. Only DRG cells with clearly visible nuclei were quantified to prevent double-counting of the same cells in different sections. For quantification, 3–5 DRG sections from 3 mice were analyzed in each group.

### Tail flick test

The ablation efficiency of the TRPV1^+^ neurons were determined by the tail flick test. The tail flick response to 51°C hot water was recorded. 20 s was set as the cut-off time.

### Chemical ablation of Trpv1^+^ and MrgprA3^+^ neurons

For systemic DTX-mediated cell ablation, *Trpv1^Cre^*; *iDTR* and *MrgprA3^GFP-Cre^; iDTR* mice were treated with diphtheria toxin (DTX), according to the previous methods with a little modification (*8*). Briefly, 6 weeks old mice were injected intraperitoneally with DTX (50 mg/kg; Sigma-Aldrich, St. Louis, MO) in 100 µl PBS at day 1, day 4, day7 and day 10. Mice were allowed for four weeks rest before behavioral experiments. For chemical ablation of Trpv1^+^ neurons, C57BL/6 mice at 4 weeks of age were treated with resiniferatoxin (RTX, Tocris Bioscience, USA). RTX was prepared in 2% DMSO with 0.15% Tween 80 in PBS. Mice received subcutaneous injections of RTX in the flank on consecutive days with three increasing doses of RTX (30, 70, and 100 μg/kg). We performed behavioral experiments 4 to 6 weeks after RTX injection.

### Mouse DRG neuron cultures

Mice were deeply anaesthetized with isoflurane and then exsanguinated. The spinal cord was removed and DRG from thoracic levels were dissected. Isolated ganglia were collected in ice-cold Ca^2+^ and Mg^2+^-free Hank’s buffered salt solution (HBSS, Gibco, USA). DRG neurons were enzymatically digested in dispase (5mg/mL, Gibco, USA) and collagenase type I (1mg/mL, Gibco, USA) dissolved in HBSS for 30 min at 37LJ, as described previously with a little modification^7^. After digestion, neurons were pelleted, suspended in neurobasal medium containing 2% B-27 supplement, 1% L-glutamine, 100 U·mL^−1^ penicillin plus 100 μg·mL^−1^ streptomycin, and 50ng·mL^−1^ nerve growth factor, plated on a 5mm coverslip coated with poly-l-lysine (10 μg·mL^−1^) and cultured under a humidified atmosphere of 5% CO2/95% air at 37°C for 18–24 h before use.

### Whole-cell patch clamp recordings

Whole-cell patch-clamp recordings were performed using a Multiclamp 700B amplifier (Molecular Devices, Sunnyvale, CA, USA) at room temperature (22–24°C) on the stage of an inverted phasecontrast microscope equipped with a filter set for tdTomato visualization. Pipettes pulled from borosilicate glass (BF 150-86-10; Sutter Instrument Company, Novato, CA, USA) with a Sutter P-1000 pipette puller had resistances of 2–4 MΩ when filled with pipette solution containing 120 mM K^+^-gluconate, 30 mM KCl, 2 mM MgCl_2_, 1 mM CaCl_2_, 2 mM MgATP, 11 mM EGTA, and 10 mM HEPES with pH 7.3. Cells were continuously perfused with standard extracellular solution containing 145 mM NaCl, 3 mM KCl, 2 mM CaCl_2_, 2 mM MgCl_2_, 10 mM glucose and 10 mM HEPES (pH was adjusted to 7.4 with NaOH). Cells were clamped to a holding potential of −80 mV and stimulated with a series of mechanical stimuli in 1 μm increments every 10 s, and the stimulus was applied for 100 ms. Step indentations were applied using a fire-polished glass pipette (tip diameter 2–3 μm) that was positioned at an angle of 70° to the surface of the cell. The pipette was controlled by a Piezo Servo Controller (E-625, Physik Instrument, Karlsruhe, Germany). Data were acquired using pClampex 10 (Molecular Devices, San Jose, CA). Currents were filtered at 2 kHz and digitized at 10 kHz. Values were given as mean ± SEM; n represents the number of measurements.

### Statistical analysis

Results are expressed as mean ± SEM. All histology, calcium imaging, and electrophysiology experiments were repeated using tissues from at least three different mice. Statistical analysis was performed in Prism 7 (GraphPad). A threshold of p < 0.05 was accepted as statistically different and p > 0.05 considered non-significant. For itch behavioral assessment, data were subjected to unpaired two-tailed Student’s *t* test (for two groups) or one-way ANOVA (for three or more groups). For statistical analysis of incidence of electrophysiological results, data were analyzed with Chisquare test.

## SUPPLEMENTAL INFORMATION

**Supplementary Figure 1.**
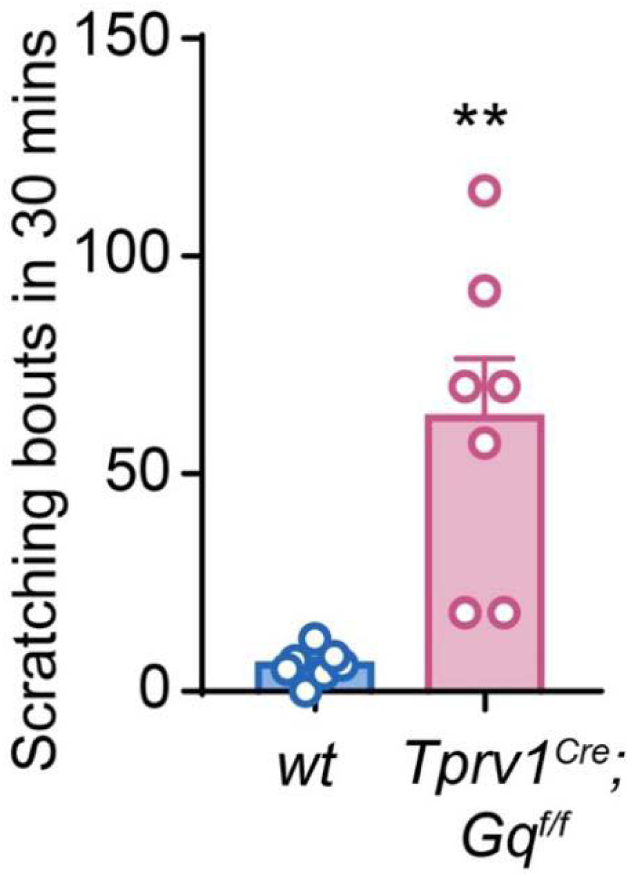
Chemogenetic activation of TRPV1+ neurons evoked spontaneous scratching behavior in the Trpv1-hM3Dq mice. n=7 for each group. Data was shown as mean ± SEM. **P<0.01. Unpaired two-tailed Student’s t test.

**Supplementary Figure 2.**
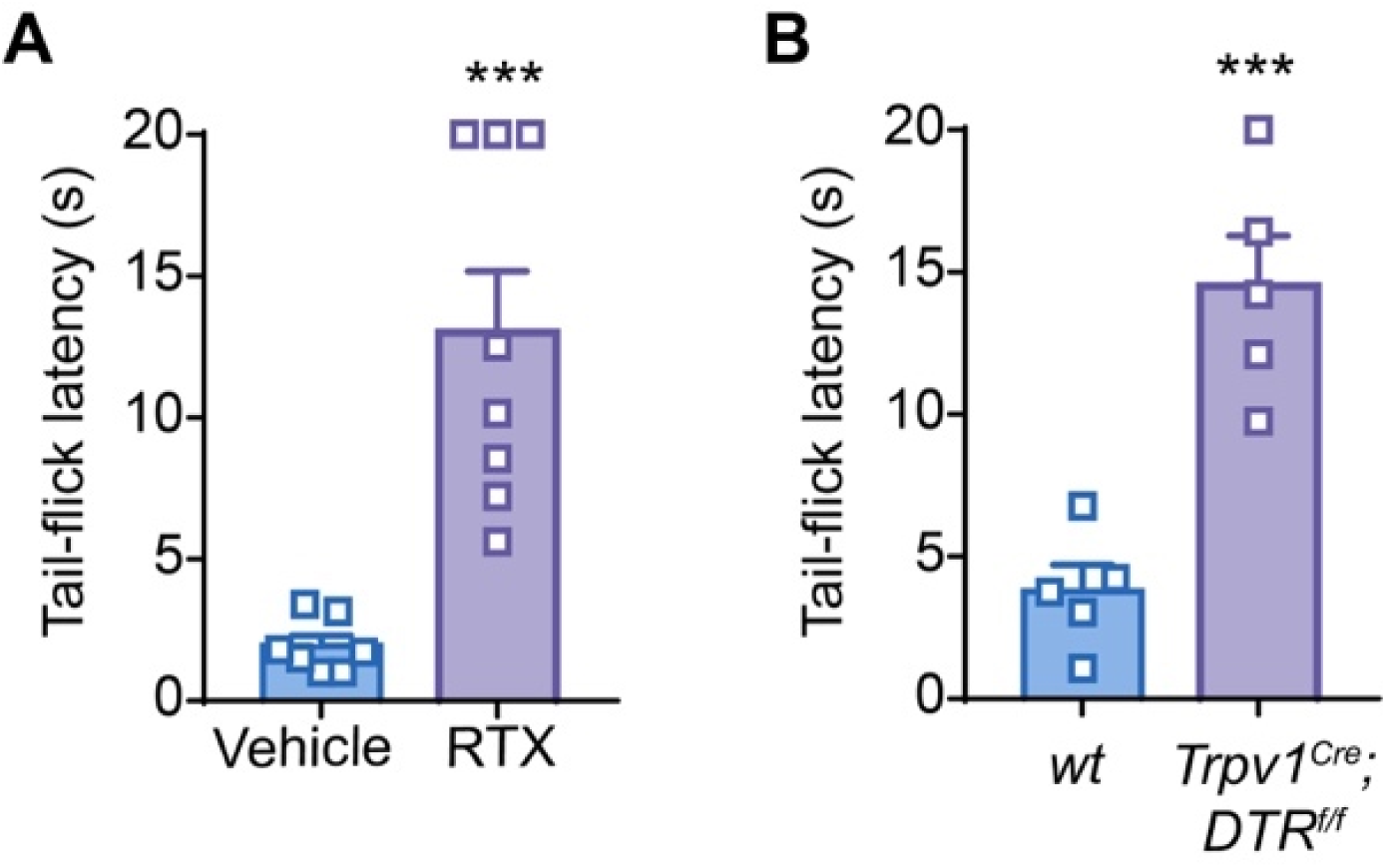
Validation of the efficiency of the TRPV1^+^ neuron ablation. **A,** The tail-flick latency of *wt* mice treated with vehicle (n=7) or RTX (n=8). **B**, The tail-flick latency of the *Cre^-^* (n=5) and *Cre^+^* (n=5) *TRPV1^Cre^*; *DTR^f/f^* mice treated with DTX. Data was shown as mean ± SEM. \*\*\**P*<0.001. Unpaired two-tailed Student’s t test.

**Supplementary Figure 3.**
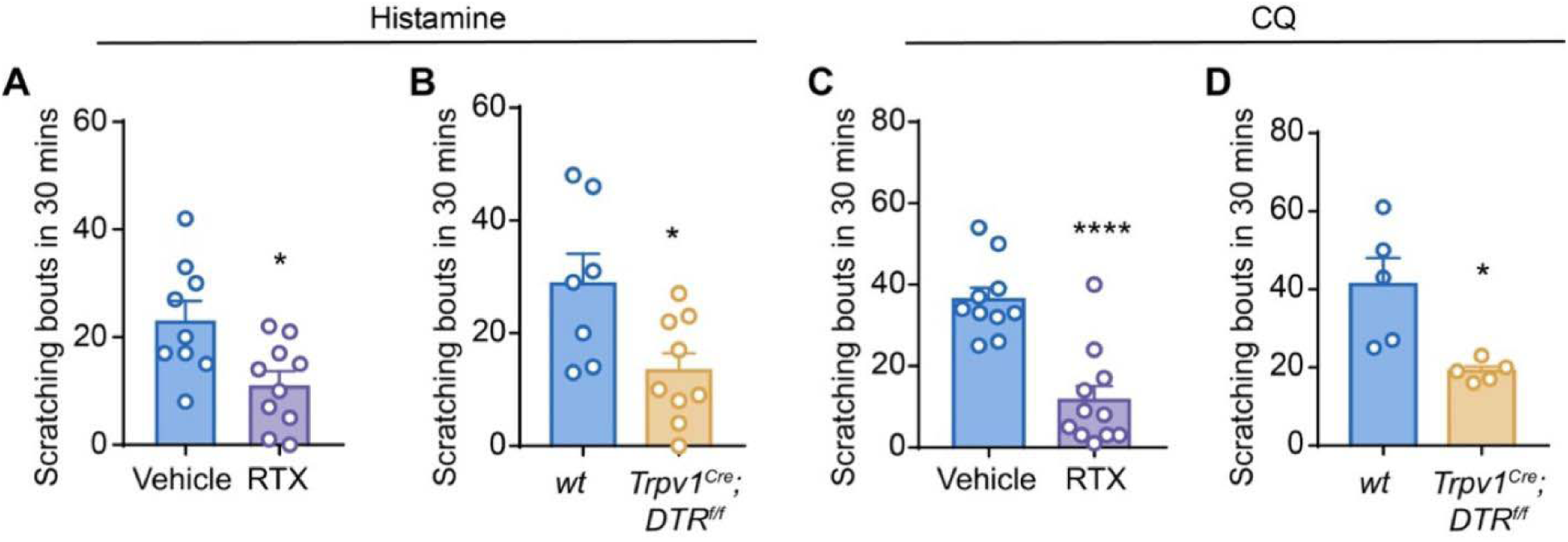
Pharmacological ablation of TRPV1^+^ neurons reduced pruritogens-induced acute itch. **A,** Intradermal injection of histamine induced acute itch in RTX-treated mice (n=10) and vehicle-treated littermates (n=9). **B**, Intradermal injection of histamine induced acute itch in the *Cre^-^*(n=7) and *Cre^+^* (n=9) *Trpv1-DTR* mice. **C**, Intradermal injection of CQ induced acute itch in RTX-treated mice (n=11) and vehicle-treated littermates (n=10). **D**, Intradermal injection of CQ induced acute itch in the *Cre^-^* (n=5) and *Cre^+^* (n=5) *Trpv1-DTR* mice. Data was shown as mean ± SEM. n.s, not significant. \**P*<0.05, \*\*\*\**P*<0.0001. Unpaired two-tailed Student’s t test.

**Supplementary Figure 4.**
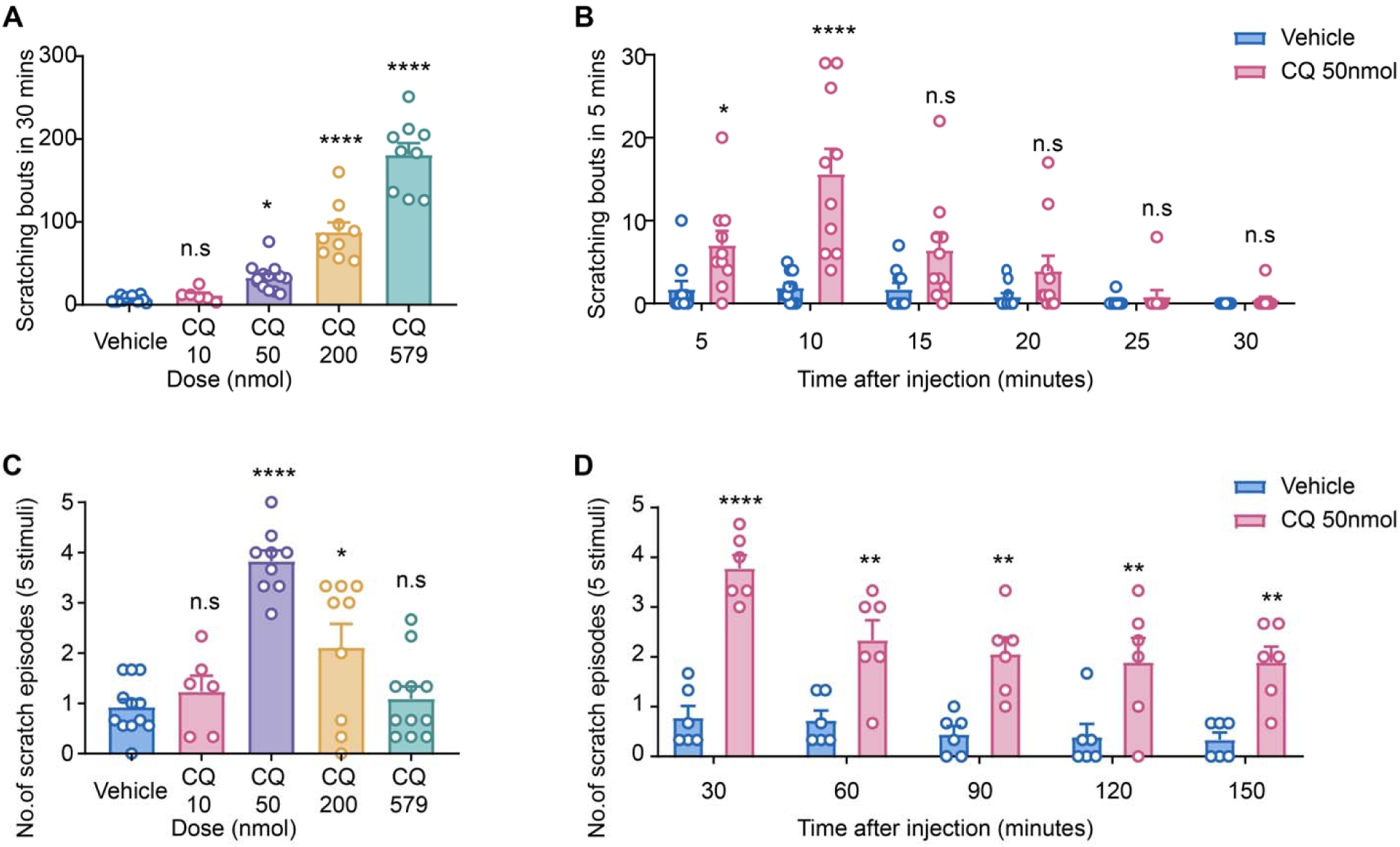
Establishment of CQ-induced mechanical itch model in mice. **A,** Dose dependency of acute itch evoked by CQ. **B,** Time course of 50 nmol CQ-induced acute itch. **C**, Dose dependency of mechanical itch evoked by CQ. **D**, Time course of 50 nmol CQ-induced mechanical itch. Data shown as mean ± SEM. n.s, not significant. \**P*<0.05, \*\**P*<0.01, \*\*\*\**P*<0.0001. One way ANOVA in A and C and unpaired two-tailed Student’s t test in B and D.

**Supplementary Figure 5.**
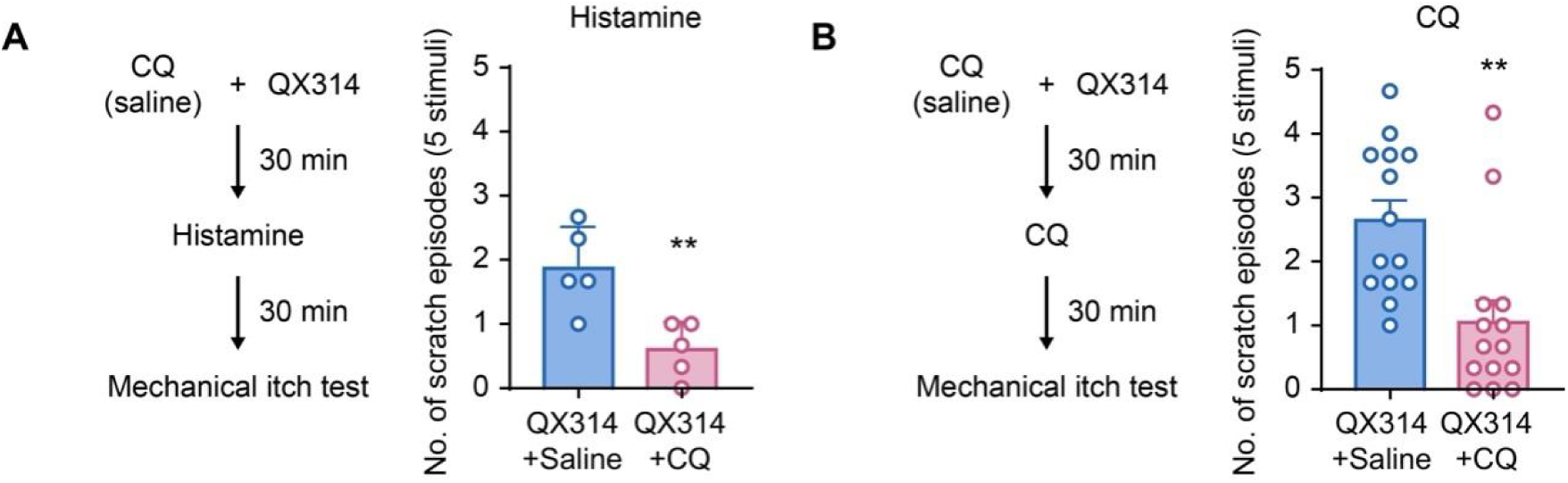
Pharmacological silencing of TRPV1^+^ neurons reduced pruritogens-induced mechanical itch. **A,** Histamine-induced and CQ-induced (B) mechanical itch in mice treated with QX314 + vehicle or QX314 + CQ. n=5 for each group. **B**, CQ-induced mechanical itch in mice treated with QX314 + vehicle or QX314 + CQ. n=14 for each group in B. Data was shown as mean ± SEM. n.s, not significant. \*\**P*<0.01. Unpaired two-tailed Student’s t test.

**Supplementary Figure 6.**
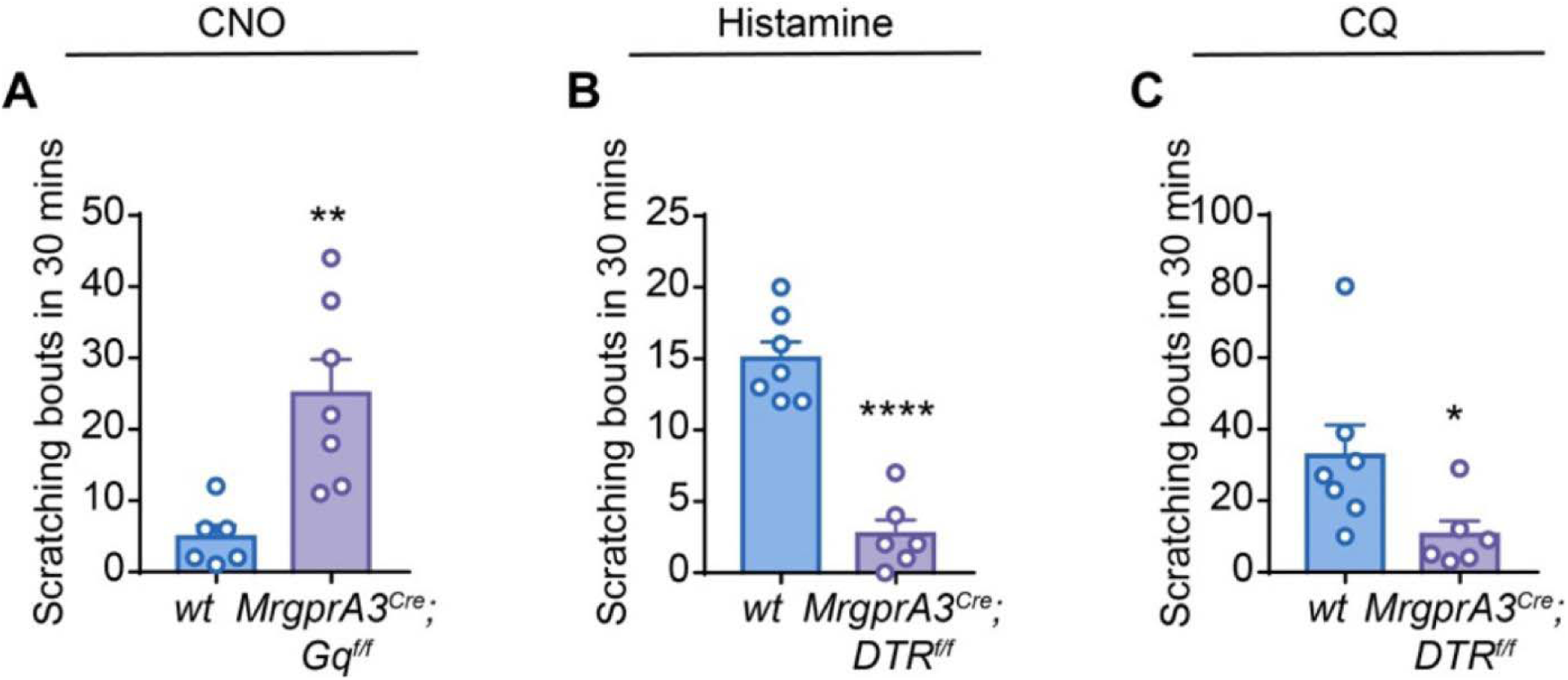
MrgprA3^+^ neurons are critically involved in the acute chemical itch. **A,** Intradermal injection of CNO evoked spontaneous scratching behavior in the *Cre^-^* (n=6) and *Cre^+^* (n=7) *MrgprA3-hM3Dq* mice. **B**, Intradermal injection of histamine evoked spontaneous scratching behavior in the *Cre^-^* (n=7) and *Cre^+^*(n=6) *MrgprA3-DTR* mice. **C**, Intradermal injection of CQ evoked spontaneous scratching behavior in the *Cre^-^* (n=7) and *Cre^+^*(n=6) *MrgprA3-DTR* mice. Data was shown as mean ± SEM. n.s, not significant. \**P*<0.05, \*\**P*<0.01, \*\*\**P*<0.001. Unpaired two-tailed Student’s t test.

**Supplementary Figure 7.**
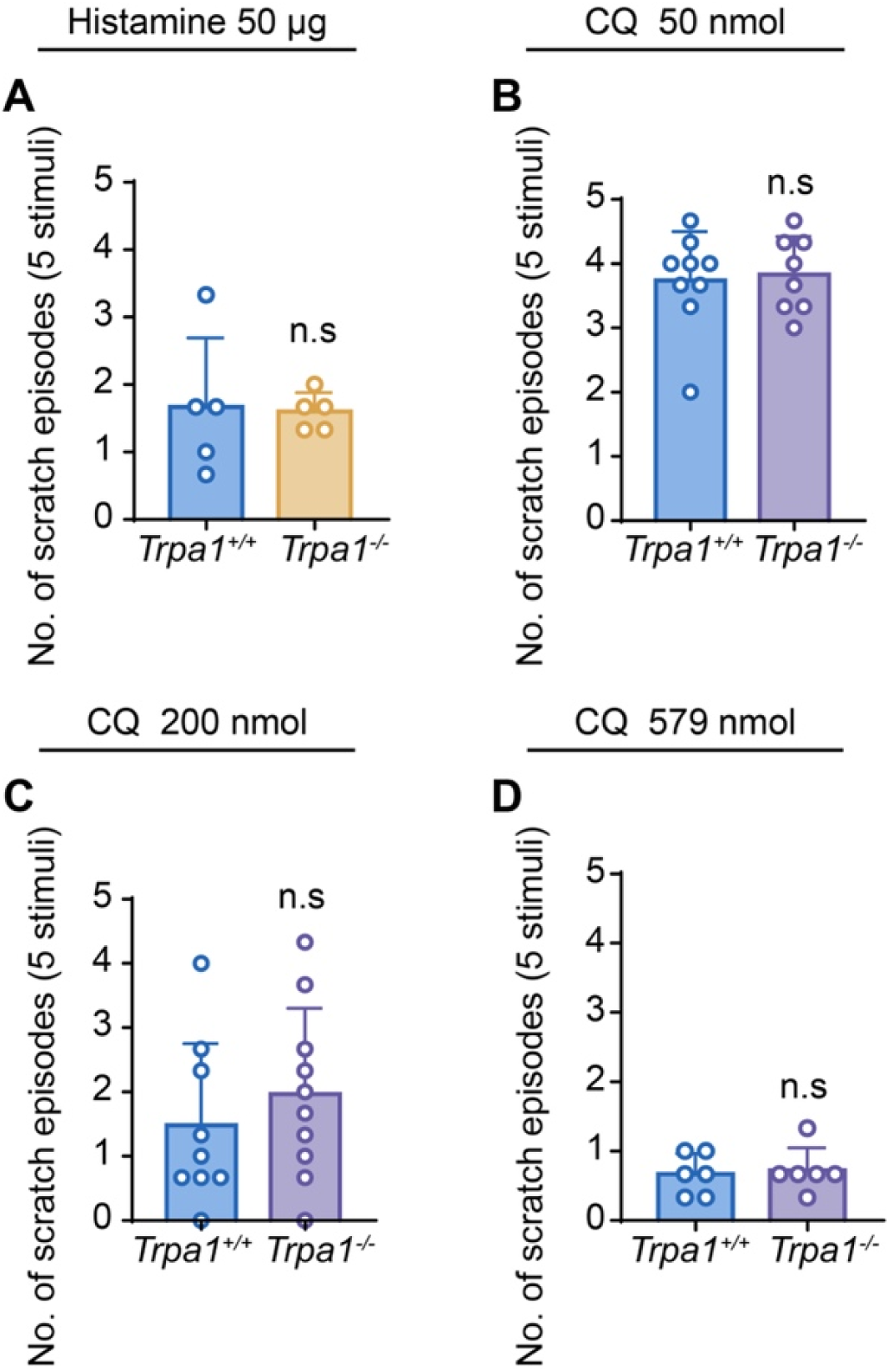
Trpa1 is not required for Histamine- and CQ-induced mechanical itch. **A,** Histamine induced alloknesis in *wt* (n=5) and *Trpa1^-/-^* mice (n=5). **B** to **D**, Different dose of CQ-induced alloknesis score in *wt* and *Trpa1^-/-^* mice. For 50 nmol CQ treatment, n=9 in *wt* group, n=8 in *Trpa1^-/-^* group; For 200 nmol CQ treatment, n=9 in *wt* group, n=10 in *Trpa1^-/-^* group; For 579 nmol CQ treatment, n=6 in each group. Data shown as mean ± SEM. n.s, not significant. Unpaired two-tailed Student’s t test.

**Supplementary Figure 8.**
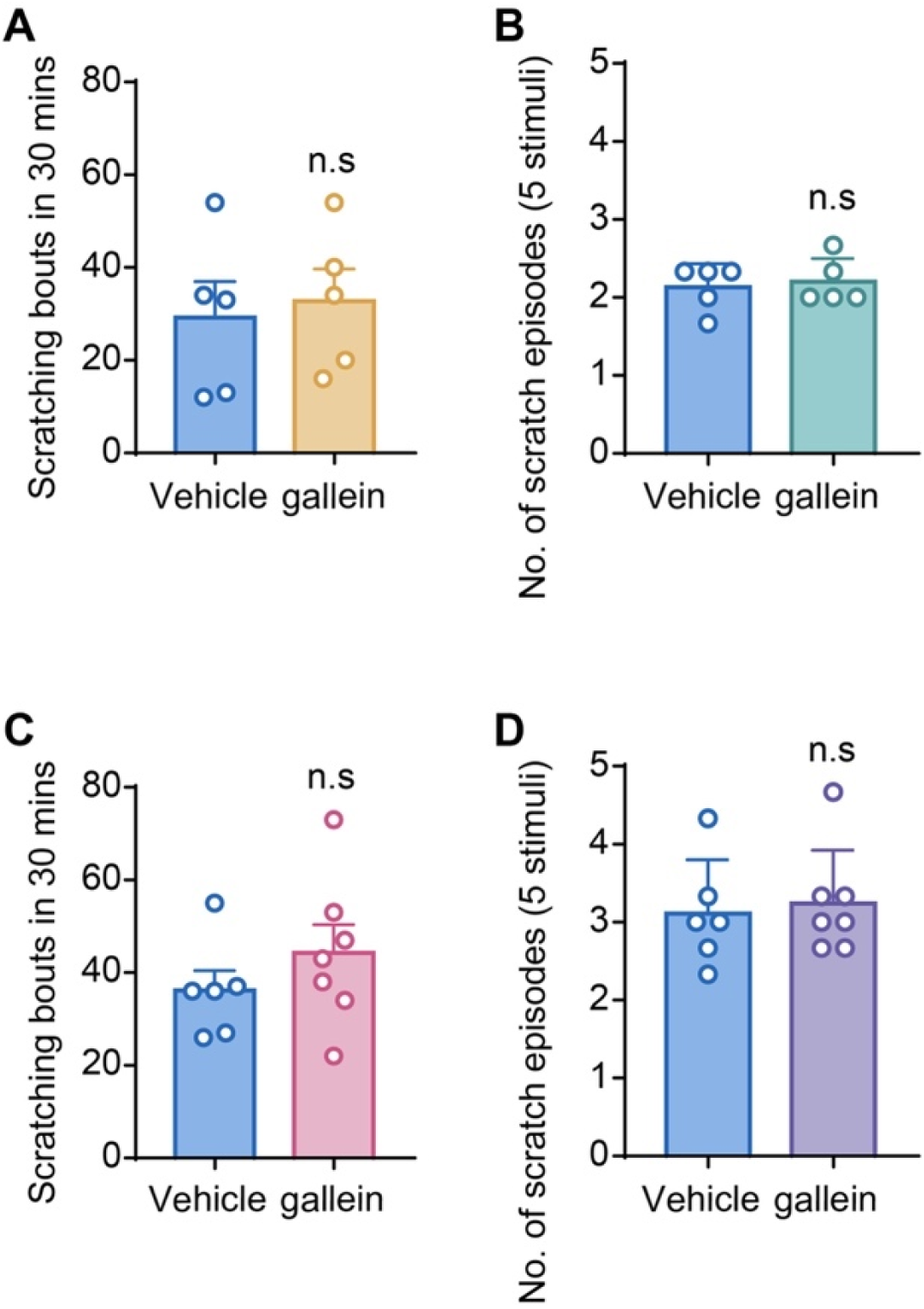
Pharmacological inhibitor of Gβγ signaling does not affect pruritogens-induced acute chemical itch or mechanical itch. **A-B,** Intradermal injection of histamine induced acute itch (A) and mechanical itch (B) in the vehicle-treated (n=5) and gallein-treated (n=5) mice. **C-D,** Intradermal injection of CQ induced acute itch (C) and mechanical itch (D) in the vehicle-treated (n=6) and gallein-treated (n=7) mice. Data was shown as mean ± SEM. n.s, not significant. Unpaired two-tailed Student’s t test.

**Supplementary Figure 9.**
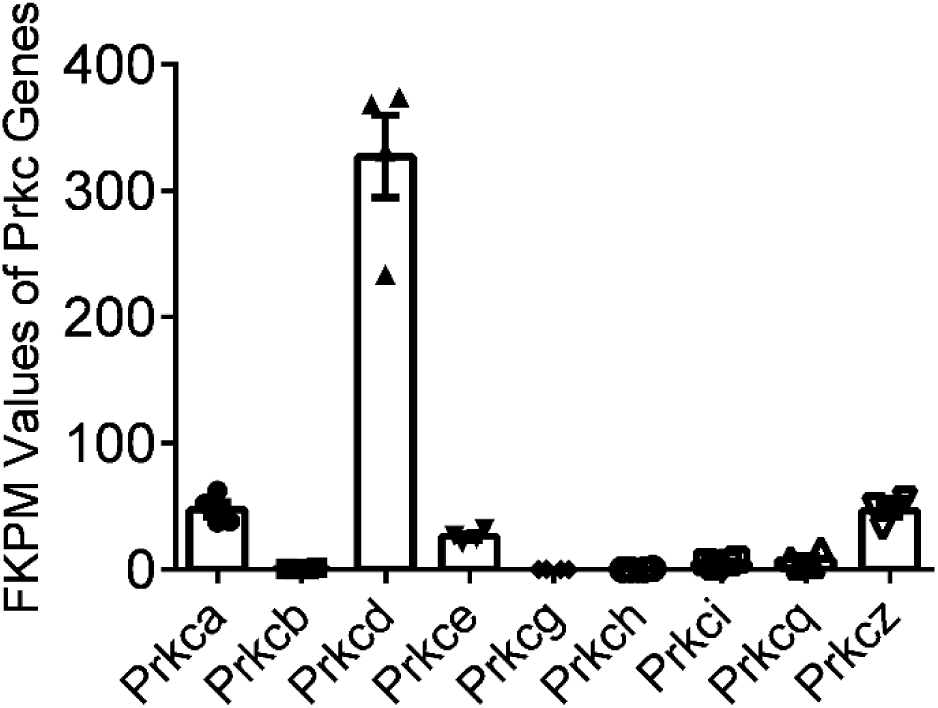
Summary data showed the expression levels (FPKM) of Prkc genes in MrgprA3^+^ neurons. Data were from a published paper by Yanyan Xing et al (Supplementary Table S5 in the original paper) and the normalized expression levels were reported as fragments per kilobase of transcript per million mapped reads (FPKM).

**Supplementary Figure 10.**
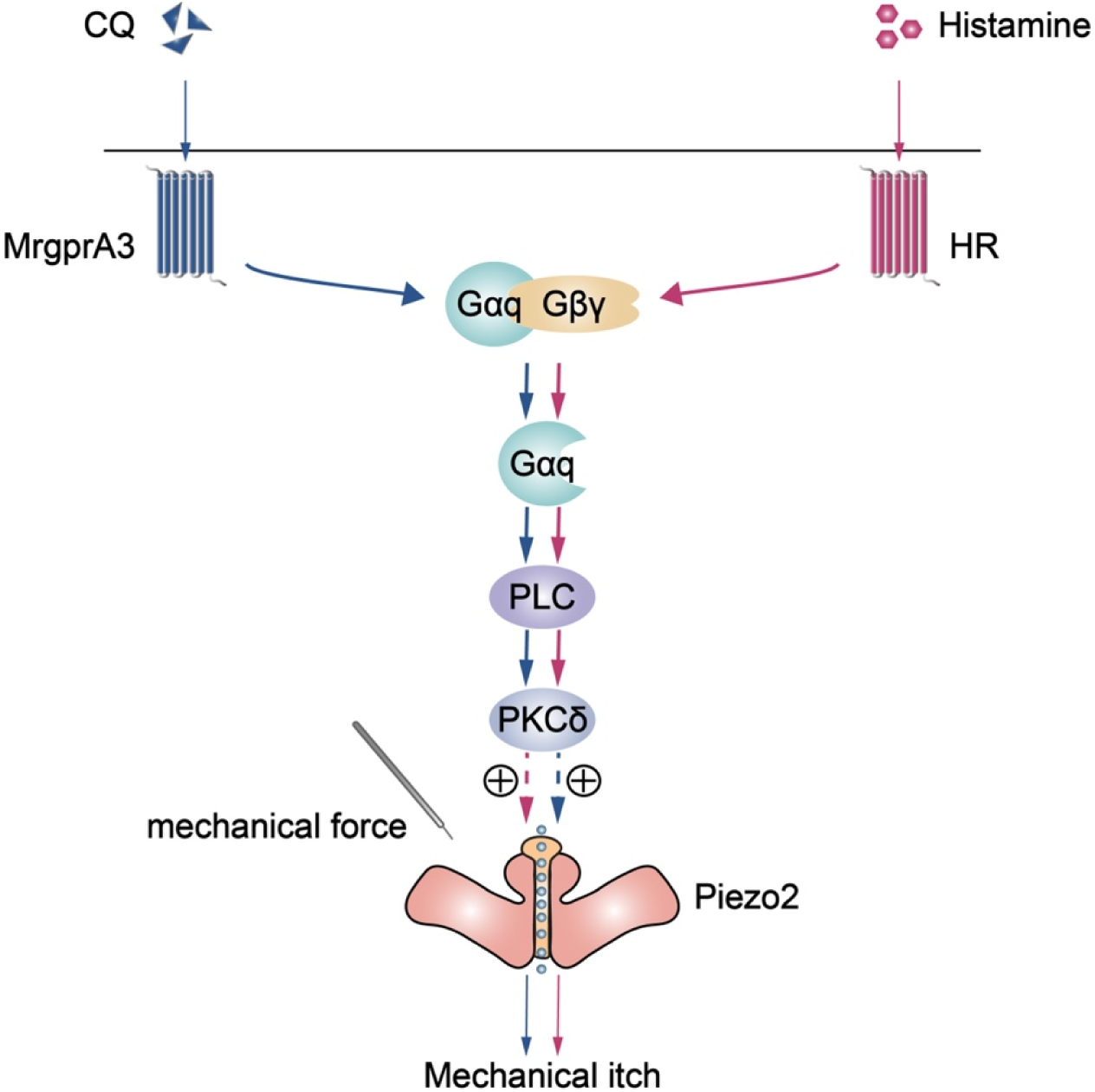
Schematic illustrating the signaling pathway in pruritogens-induced alloknesis. In response pruritogens-induced activation, GPCR-dissociated G_αq_ activates downstream PLC/PKCδ signaling, resulting in the sensitization of pruriceptor-expressing Piezo2 channel and contributing to the initiation of alloknesis.

